# The bithorax complex *iab-7* Polycomb Response Element has a novel role in the functioning of the *Fab-7* chromatin boundary

**DOI:** 10.1101/329284

**Authors:** Olga Kyrchanova, Amina Kurbidaeva, Marat Sabirov, Nikolay Postika, Daniel Wolle, Tsutomu Aoki, Oksana Maksimenko, Vladic Mogila, Paul Schedl, Pavel Georgiev

## Abstract

Expression of the three *Bithorax* complex homeotic genes is orchestrated by nine parasegment-specific regulatory domains. Autonomy of each domain is conferred by boundary elements (insulators). Here, we have used an *in situ* replacement strategy to reanalyze the sequences required for the functioning of one of the best-characterized fly boundaries, *Fab-7*. It was initially identified by a deletion, *Fab-7*^*1*^, that transformed parasegment (PS) 11 into a duplicate copy of PS12. *Fab-7*^*1*^ deleted four nuclease hypersensitive sites, HS*, HS1, HS2, and HS3, located in between the *iab-6* and *iab-7* regulatory domains. Transgene and *P*-element excision experiments mapped the boundary to HS*+HS1+HS2, while HS3 was shown to be the *iab-7* Polycomb response element (PRE). Recent replacement experiments showed that HS1 is both necessary and sufficient for boundary activity when HS3 is also presented in the replacement construct. Surprisingly, while HS1+HS3 combination has full boundary activity, we discovered that HS1 alone has only minimal function. Moreover, when combined with HS3, only the distal half of HS1, dHS1, is needed. A ∼1,000 kD multiprotein complex containing the GAF protein, called the LBC, binds to the dHS1 sequence and we show that mutations in dHS1 that disrupt LBC binding in nuclear extracts eliminate boundary activity and GAF binding *in vivo*. HS3 has binding sites for GAF and Pho proteins that are required for PRE silencing. In contrast, HS3 boundary activity only requires the GAF binding sites. LBC binding with HS3 in nuclear extracts, and GAF association *in vivo* depend upon the HS3 GAF sites, but not the Pho sites. Consistent with a role for the LBC in HS3 boundary activity, the boundary function of the dHS1+HS3^mPho^ combination is lost when the flies are heterozygous for a mutation in the GAF gene. Taken together, these results reveal a novel function for the *iab-7* PREs in chromosome architecture.

**Author Summary:** Polycomb group proteins (PcG) are important epigenetic regulators of developmental genes in all higher eukaryotes. In *Drosophila,* these proteins are bound to specific regulatory DNA elements called Polycomb group Response Elements (PREs). PcG support proper patterns of homeotic gene expression throughout development. *Drosophila* PREs are made up of binding sites for a complex array of DNA binding proteins, including GAF and Pho. In the regulatory region of the bithorax complex (BX-C), the boundary/insulator elements organize the autonomous regulatory domains, and their active or repressed states are regulated by PREs. Here, we studied the domain organization of the *Fab-7* boundary and the neighboring PRE, which separate the *iab-6* and *iab-7* domains involved in transcription of the *Abd-B* gene. It was previously thought that PRE recruits PcG proteins that inhibit activation of the *iab-7* enhancers in the inappropriate domains. However, here we found that PRE contributes to boundary activity and in combination with a key 242 bp *Fab-7* region (dHS1) can form a completely functional boundary. Late Boundary Complex (LBC) binds not only to dHS1 but also to PRE and is required for the boundary activity of both elements. At the same time, mutations of Pho binding sites strongly diminish recruiting of PcG but do not considerably affect boundary function, suggesting that these activities can be separated in PRE.

## Introduction

Chromosomes in multicellular organisms are subdivided into a series of independent topologically associating domains (or TADs) [1,2]. The average length of these domains in humans is 180 kb, while they are only on the order 5-20 kb in flies [3-5]. In mammals, TADs are frequently defined by binding sites for the conserved zinc finger protein CTCF [6,7]. While a single CTCF is thought to be necessary and sufficient for boundary function in mammals, this is not true in flies. More than a dozen DNA binding proteins in flies that function as architectural factors have been identified and it is likely that many more remain to be discovered [8-10]. Because of extensive redundancy any one of these individual recognition sequences for these factors might not be necessary for boundary function.

One of the best examples of functional redundancy is the *Fab-7* boundary in the *Drosophila* bithorax complex (BX-C). BX-C contains three homeotic genes, *Ultrabithorax* (*Ubx*), *abdominal-A* (*abd-A*), and *Abdominal-B* (*Abd-B*), which are responsible for specifying the parasegments (PS5 to PS13) that make up the posterior two-thirds of the fly segments [11-14]. Expression of the homeotic genes in the appropriate parasegment-specific pattern is orchestrated by a series of nine *cis-*regulatory domains, *abx/bx, bxd/pbx, iab-2— iab-9* (Figure 1A). For example, the *iab-5, iab-6, iab-7,* and *iab-8 cis*-regulatory domains direct *Abd-B* expression in PS10-PS13 [15,16]. BX-C regulation is divided into two phases, initiation and maintenance [11,17]. During the initiation phase, a combination of gap and pair-rule proteins interact with initiation elements in each regulatory domain, setting it in the *on* or *off* state. In PS10, for example, initiators in *iab-5* activate the domain, while *iab-6, iab-7,* and *iab-8* are set in the *off* state. In PS11, *iab-6* is activated, while *iab-7* and *iab-8* are *off*. Once the gap and pair-rule gene proteins disappear during gastrulation, the *on* and *off* states of the regulatory domains are maintained by Trithorax (Trx) and Polycomb (PcG) group proteins, respectively [18-21]. These maintenance factors are recruited to the domains by special cis-acting elements called Trithorax Response Elements (TREs) and Polycomb Response Elements (PREs) [22-26]. In addition to elements that establish and maintain the *on/off* state, each domain has a series of tissue and cell type specific enhancers that direct the expression of the target homeotic gene in an appropriate pattern [11,16]. For example, the tissue/cell type enhancers in *iab-6* drive *Abd-B* expression in a pattern that orchestrates the proper differentiation of cells with a PS11 identity. This pattern of expression is distinct from that in PS12, where *Abd-B* is regulated by enhancers in *iab-7*.

**Fig. 1.**
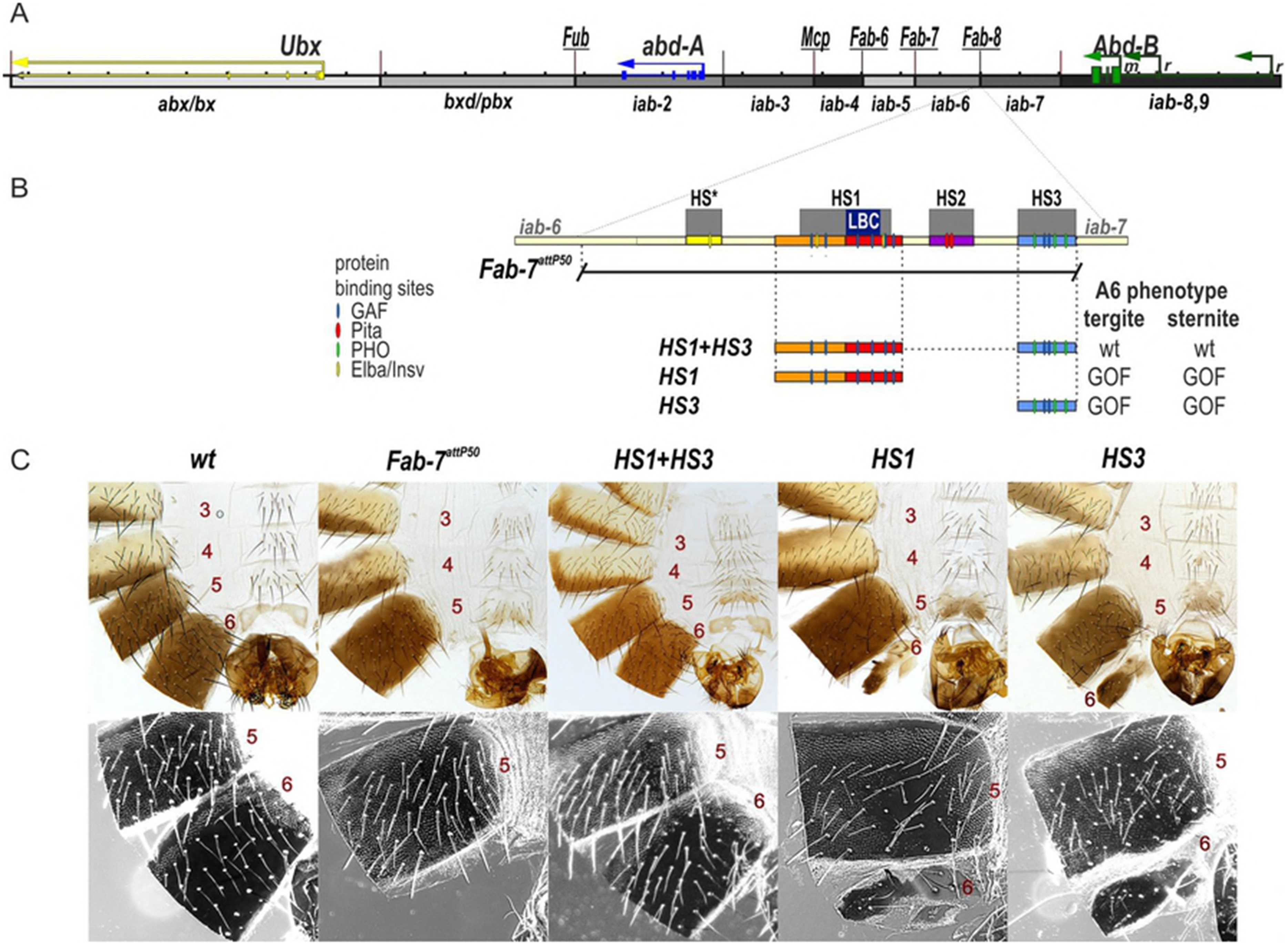
HS1 alone is not sufficient for boundary function. (A) Map of the bithorax complex showing the location of the three homeotic genes and the parasegment-specific regulatory domains. (B) Map of *Fab-7* region showing the four hypersensitive sites, HS*, HS1, HS2, and HS3. The locations of recognition motifs for proteins known to be associated with *Fab-7* are indicated. Replacement fragments are shown below the map, with a summary of their cuticle (tergite and sternite) phenotypes. (C) Bright field (top) and dark field (bottom) images of cuticles prepared from wild type (*wt*), *Fab-7* attP (replacement platform), *HS1+HS3, HS1,* and *HS3* male flies. As described in the text, the *HS1+HS3* replacement resembles wild type. In contrast, the *HS1* and *HS3* replacements have a strong GOF transformation. In those instances in which residual A6 (PS11) cuticle is present, it shows evidence of a LOF transformation.

The *Fab-7* boundary, like other boundary elements in BX-C, is required to ensure that the flanking regulatory domains, *iab-6* and *iab-7*, are able to function autonomously [27-29]. During the initiation phase it blocks crosstalk between initiators in *iab-6* and *iab-7*, while during the maintenance phase it keeps the domains in *on* or *off* state by preventing interactions between their PREs and TREs. In addition, like most other BX-C boundaries, *Fab-7* must also facilitate “bypass”, enabling the distal regulatory domains *iab-5* and *iab-6* to “jump over” and contact *Abd-B* in PS10 and PS11, respectively. Like blocking, bypass activity is essential for proper *Abd-B* regulation. *Fab-7* was initially defined by a 4 kb X-ray induced deletion that had an unusual dominant gain-of-function (GOF) phenotype, transforming PS11 into a duplicate copy of PS12 [27]. The *Fab-7*^*1*^ deletion spanned four nuclease hypersensitive regions, HS*, HS1, HS2, and HS3 [30]. Subsequent transgene studies showed that a 1.2 kb fragment spanning HS*+HS1+HS2 had enhancer blocking activity in embryos and in adults [31,32]. The fourth hypersensitive region, HS3, had no detectable boundary activity; however, when included in a *white* transgene, it induced pairing sensitive silencing, which is a characteristic activity of PREs [33]. This separation of functions was confirmed by Mihaly et al [29] who generated a series of new deletions that removed either HS*+HS1+HS2 or HS3. Unlike *Fab-7*^*1*^, deletions that removed only HS*+HS1+HS2 have a mixed GOF and LOF (loss-of-function phenotype) (PS11➔PS12 and PS1l➔P10, respectively). Mutations in PcG genes enhance the GOF phenotypes, while mutations in Trx enhance the LOF phenotypes. By contrast, PcG and Trx mutations have no effect on the GOF phenotypes of deletions, like *Fab-7*^*1*^, that remove all four hypersensitive regions. Finally, flies carrying HS3 deletions are typically wild type as heterozygotes or homozygotes and only infrequently weak GOF phenotypes are observed in homozygous animals [29,34]. While the LOF phenotypes in HS*+HS1+HS2 deletions require silencing of the *iab-6* domain by the *iab-7* PRE, this is not the only mechanism that can give rise to LOF transformations of PS11 (and PS10). LOF phenotypes are also observed when *Fab-7* is replaced by heterologous boundaries such *scs, su(Hw),* or the BX-C boundary *Mcp* [35-37]. Like *Fab-7*, these boundaries prevent crosstalk between *iab-6* and *iab-7*; however, they fail to support bypass, and instead block the *iab-6* domain from regulating *Abd-B*. Thus far, the only heterologous boundary that recapitulates both the blocking and bypass activity of *Fab-7* is the neighboring boundary *Fab-8* [36,38].

Previous studies indicate that *Fab-7* (HS*+HS1+HS2) boundary activity is generated by a combination of ubiquitously expressed factors and stage/cell type specific factors. One ubiquitously expressed factor is the zinc finger protein Pita which binds to sites in HS2 [37,39]. The known developmentally regulated factors are Elba, Insensitive (Insv), and the LBC [40]. The LBC is a multiprotein complex that contains at least three distinct DNA binding proteins, the GAGA factor (GAF), Clamp, and Mod(mdg4) [41]. In the case of *Fab-7*, three contiguous sequences spanning GAGA sites 3, 4, and 5 generate LBC shifts [34,42].

While transgene experiments indicated that sequences spanning HS*+HS1+HS2 are required for blocking activity [43,44], boundary replacement using an *attP* platform that deletes HS*+HS1+HS2+HS3 suggested that the requirements for boundary function out of context are more demanding than those in BX-C. These replacement experiments indicated that HS* and HS2 are not required for boundary activity, while the largest hypersensitive region, HS1, is both necessary and sufficient [34]. However, there was a confounding factor in these experiments: because the HS3 *iab-7* PRE has Polycomb silencing activity and as such is an important regulatory component of the *iab-7* regulatory domain, it was retained in these replacement experiments. Here we have asked whether HS1 alone is sufficient in the absence of HS3 *iab-7* PRE. Surprisingly, it is not. Instead, our studies indicate that HS3 not only has PRE activity, but also that it contributes to the boundary function of *Fab-7*. Moreover, it appears that like HS1, the LBC is important for the boundary (and PRE) functions of HS3.

## Results

### HS3 rescues boundary activity of HS1

Studies by Wolle et al [34] have shown that HS1 is both necessary and sufficient for boundary function. However, in these experiments HS3 was also present. For this reason we wondered whether HS1 would be sufficient in the absence of HS3. Figure 1B shows the *HS1* replacement, and as controls, the starting platform *Fab-7*^*attP50*^, *HS1+HS3,* and *HS3* alone. The *Fab-7*^*attP50*^ platform deletes *HS*+HS1+HS2+HS3*, and as observed for the *Fab-7*^*1*^ deletion, the A6 segment is completely transformed into a duplicate copy of A7 (Figure 1C). As reported previously, we found that flies carrying the *HS1+HS3* replacement are indistinguishable from wild type, while for the *HS3* replacement, there is a strong GOF transformation, and the A6 tergite is greatly reduced in size or absent, while the sternite is missing (Figures 1C and S1). Unexpectedly, like *HS3, HS1* alone also has a strong GOF phenotype. The sternite is typically missing, while the tergite is typically greatly reduced in size (Figures 1C and S1).

The fact that HS1 is insufficient for boundary function in the absence of HS3 prompted us to reassess the role of HS3 in *Fab-7* boundary function. To address this question, we examined the functional properties of different combinations of the HS*, HS1, and HS2 regions either in the presence or absence of the *iab-7* PRE, HS3. We first attempted to rebuild the boundary using combinations of HS*, HS1, and HS2. As previously reported in studies on the Pita sites in HS2, *HS1+HS2+HS3,* or *HS1+HS2*^*ΔPita*^ *+HS3* replacements are fully functional [37]. In contrast, the boundary function of the *HS1+HS2* replacement is tissue-specific (Figure 2). In *HS1+HS2* flies, the A6 tergite is fully wild type. In contrast, the sternite is absent indicative of a GOF transformation of PS11➔PS12 in the cells that give rise to the ventral cuticle. In this context, the Pita sites in HS2 are essential for boundary function in the cells that give rise to the tergite [37].

**Fig. 2.**
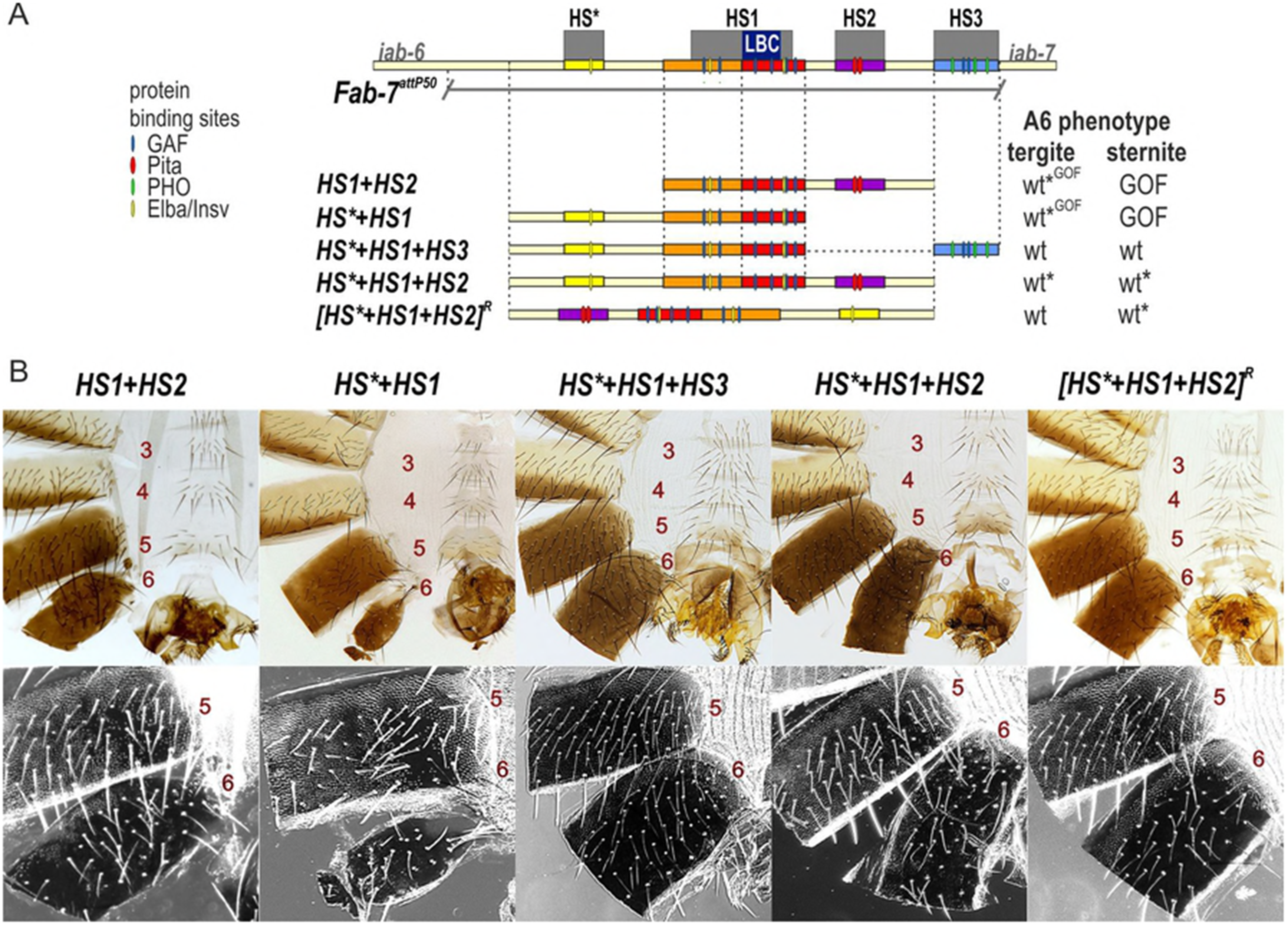
HS1 combinations with either HS2 or HS* are not fully functional. (A) Map of *Fab-7* region showing the four hypersensitive sites, HS*, HS1, HS2, and HS3 and the locations of recognition motifs for proteins known to be associated with *Fab-7*. Replacement fragments are shown below the map with a summary of their cuticle (tergite and sternite) phenotypes. (B) Bright field (top) and dark field images (bottom) of cuticles prepared from *HS1+HS2, HS*+HS1+HS3, HS*+HS1, HS*+HS1+HS2*, and *[HS*+HS1+HS2]R* (reverse) male flies. As detailed in the text, the *HS1+HS2* replacement lacks a sternite (GOF), but has a nearly normal tergite size with wild type morphology. The *HS*+HS1+HS3* looks like wild type. The *HS*+HS1* replacement lacks a sternite (GOF), while there is a variable reduction in the size of tergite. The morphology of the residual tergite suggests that it has the appropriate A6 (PS11) identity. *HS*+HS1+HS2* males frequently show some A6 cuticle defects indicative of a weak A6(PS11)➔A7(PS12) (also see Figure S2). *[HS*+HS1+HS2]R* flies have a normal A6 tergite; however, the sternite shows evidence of a weak LOF transformation (bristles). wt* -- minor deviations in phenotype. wt*^GOF^ – variable phenotype between wt and GOF.

We next tested the *HS*+HS1* combination with or without HS3. While the *HS*+HS1+H3* combination is fully functional, *HS*+HS1* retains only limited tissue-specific boundary activity (Figure 2). *HS*+HS2* appears to have no boundary activity in the histoblasts giving rise to the ventral cuticle, and in all male flies the A6 sternite is completely absent. In about 80% of the males, the A6 tergite is greatly reduced in size and has an irregular shape, as expected for an A6➔A7 (or PS11➔PS12) transformation in segment identity (Figures 2 and S2). In the remaining 20%, the boundary appears to be at least partially functional and there is only a slight reduction in the size of the A6 tergite.

The experiments described above indicate that HS3 complements the blocking defects of boundaries composed of just HS1 or HS1 plus either HS2 or HS*. We wondered whether HS3 might also contribute to the bypass activity. To test this possibility, we reinvestigated the effects of inverting the *Fab-7* boundary. In previous study, we showed that the blocking and bypass activity of *HS1+HS2* is orientation independent [36].

However, this experiment was done in the presence of HS3. To determine if HS3 contributes to orientation independence, we generated forward and reverse *HS*+HS1+HS2* replacements that lacked HS3 (Figure 2). The properties of the *HS*+HS1+HS2* replacement resemble Class III deletions described by Mihaly et al [29]. While the size of A6 tergite is normal in all but about 5% males, the sternites are typically thinner and slightly malformed. This phenotype is indicative of a very weak GOF transformation of the A6 sternite (Figures 2 and S2). A different result was obtained for the inverted *[HS*+HS1+HS2]*^*R*^ replacement (Figure 2). All flies had a fully wild type tergite, while the sternites were weakly malformed, exhibiting weak LOF transformation of A6 into A5. Since the HS3 *iab-7* PRE was absent in these replacements, the weak LOF phenotypes are expected to arise because bypass activity is partially compromised in cells giving rise to the ventral adult cuticle.

### Functional dissection of HS1

In contrast to HS1 alone, the *HS1+HS3* combination has full boundary function. To further probe the requirements for HS1 boundary activity when combined with HS3, we subdivided HS1 into proximal and distal parts, pHS1 and dHS1 (Figure 3). Previous experiments have implicated the LBC in the late blocking activity of dHS1, while an Elba/Insv recognition sequence is likely to contribute to early blocking activity of pHS1 [40]. Figures 3 and S3 show that the *pHS1+HS3* combination gives a strong GOF transformation of A6, indicating that the pHS1 sequence is not able to reconstitute boundary activity. In contrast, the *dHS1+HS3* combination has a fully wild type A6 segment, just like the *HS1+HS3* combination.

**Fig. 3.**
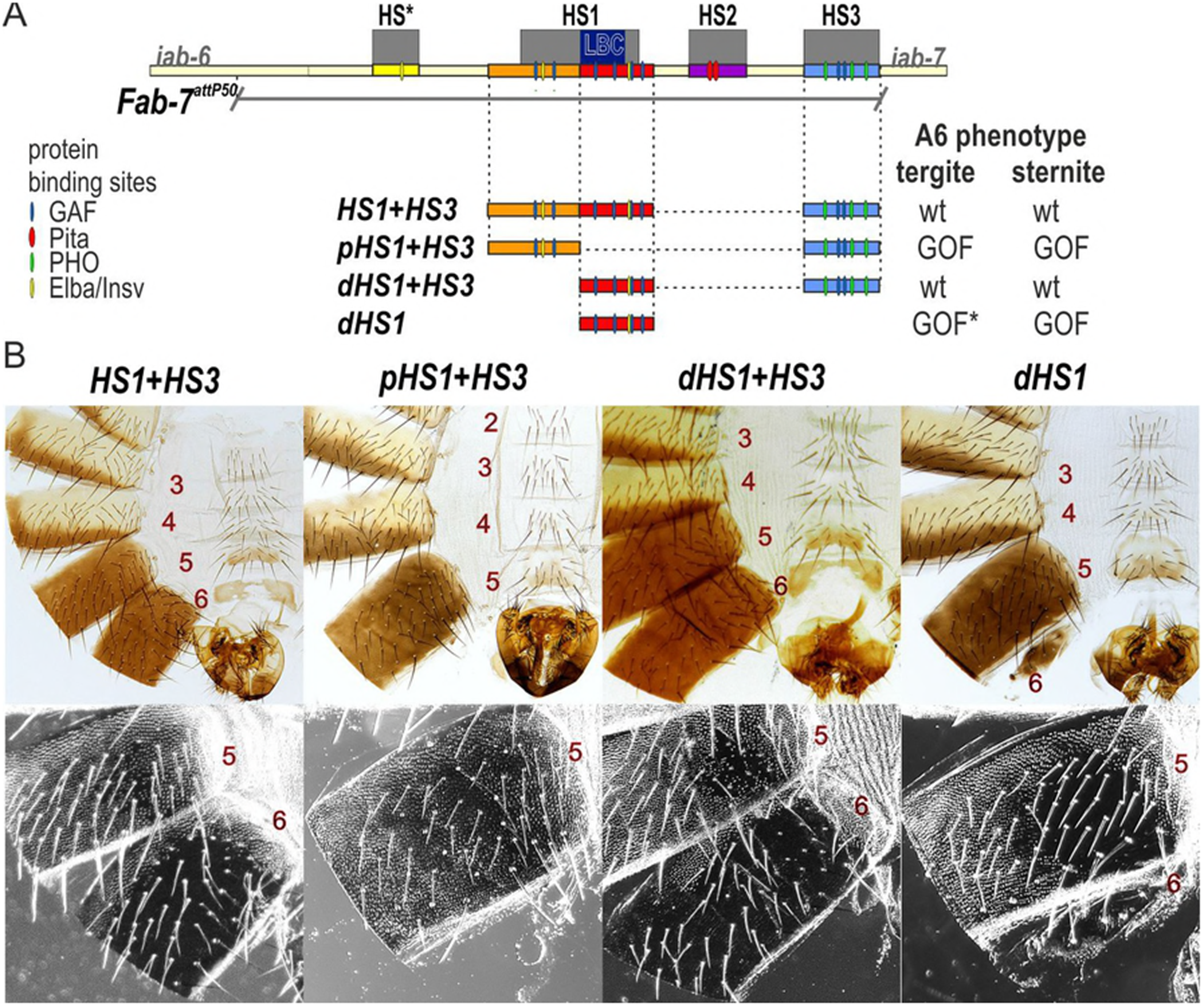
dHS1 but not pHS1 can function as a boundary together with HS3. (A) Map of *Fab-7* region showing the four hypersensitive sites, HS*, HS1, HS2, and HS3 and the location of recognition motifs for proteins known to be associated with *Fab-7*. Replacement fragments are shown below the map with a summary of their cuticle (tergite and sternite) phenotypes. (B) Bright field (top) and dark field images of cuticles prepared from *HS1+HS3, pHS1+HS3, dHS1+HS3,* and *dHS1* male flies. As described in text, the *HS1+HS3* and *dHS1+HS3* replacements resemble wild type males, while *pHS1+HS3* and *dHS1* flies have strong GOF phenotypes. In flies that have residual A6 cuticle (typically, a tergite), there are LOF transformations. GOF* -- incomplete GOF phenotype in most males.

Wolle et al [34] showed that the LBC can bind independently to ∼65-80 bp probes spanning the GAGA3, GAGA4, and GAGA5 sequences in dHS1, and that binding to the 65 bp GAGA3 and GAGA4 probes requires the GAGAG motif. However, we subsequently found that optimal LBC binding is to larger DNA probes that span GAGA3-4, GAGA3-5, or even GAGA3-6. The experiments shown in Figure S4 A, B compare LBC binding to the 65 bp GAGA3 and GAGA4 probes with binding to a larger GAGA3+4 probe. In both binding and competition experiments we found that there is a 6-12 fold differential in LBC binding to the larger GAGA3+4 probe. This difference in relative affinity isn’t due to just probe length. The competition experiment in Figure S4C shows that a hybrid GAGA3+LacZ probe of the same length as GAGA3+4 is a poor competitor for LBC binding compared to GAGA3+4. Even larger differences in relative affinity are observed for probes spanning GAGA3-5 or GAGA3-6.

While LBC binding to the 65 bp GAGA3 and GAGA4 probes requires the GAGAG motif, how mutations in this motif affect LBC binding to larger dHS1 probes hasn’t been investigated. For this purpose, we compared LBC binding to dHS1 probes that are wild type or have mutations in GAGA3-6. Figure 4A shows that LBC binding to the dHS1 probe is largely abrogated when all four GAGA motifs are mutant (GAGAmut3-6).

**Fig. 4.**
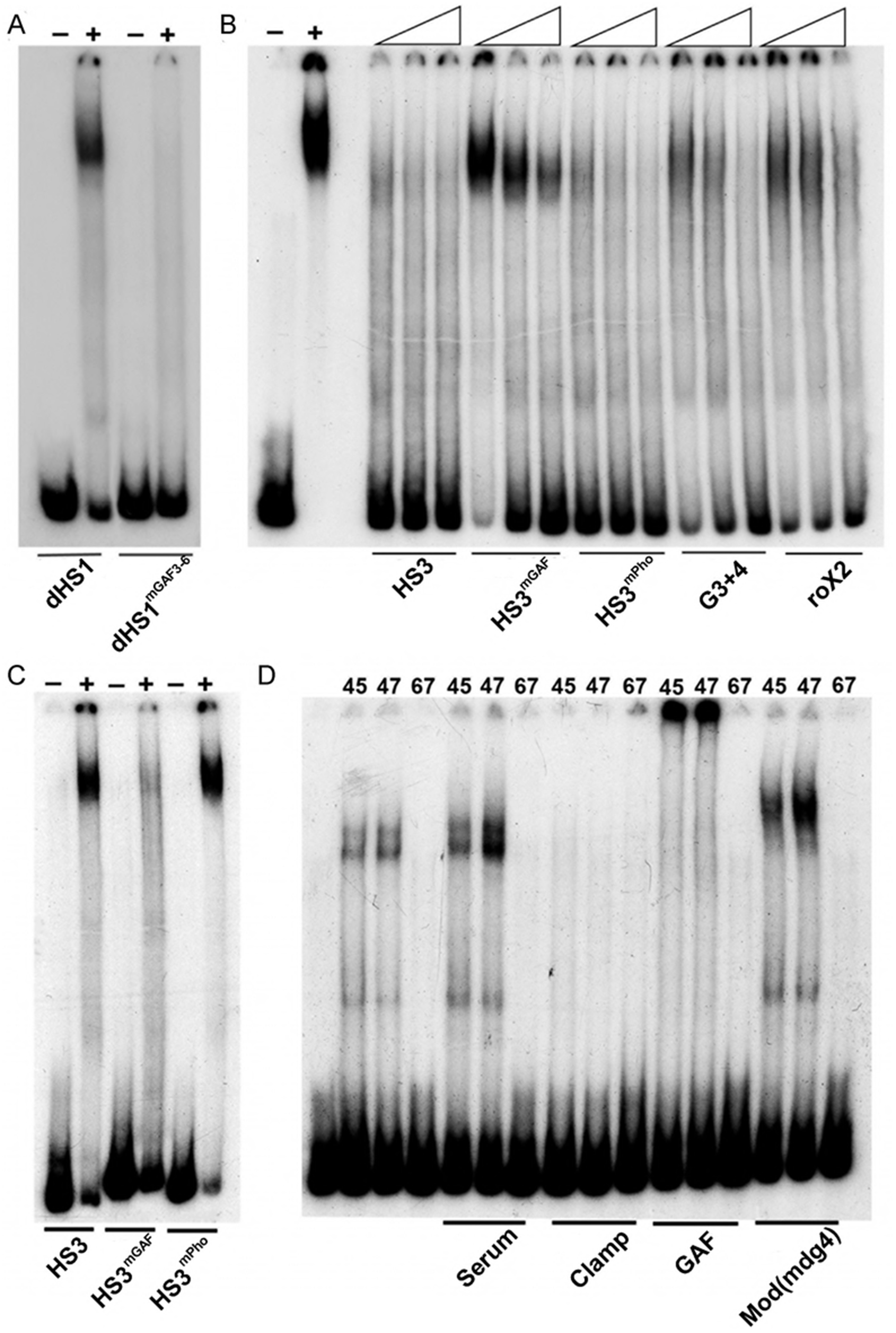
LBC binding to dHS1 and HS3 requires the GAGAG motifs. Nuclear extracts prepared from 6-18 hr embryos were used for EMSA experiments: (-) no extract, (+) with extract. (A) EMSA experiments with a wild type and GAGAG mutant (as indicated) dHS1 probe. (B) EMSA competition experiments with a probe spanning HS3. Control lanes on the left show the LBC shift of HS3 in the absence of cold competitors. As illustrated by the triangles, increasing concentrations (25x, 50x, 100x) of cold competitor were added. The cold competitor used in each set of three lanes is indicated below. (C) EMSA of wild type and mutant HS3 probes. The two HS3 GAGAG sites are mutant in the HS3^mGAF^ probe. The three HS3 Pho sites are mutant in the HS3^mP^ probe. (D) Antibody supershift experiments using fractions from a gel filtration column. Fraction numbers (45, 47, and 67) are indicated above each lane. 45 and 47 are two of LBC peak fractions while fraction 67 doesn’t have LBC activity. The antibody used for each set of supershift experiments is indicated below. Note: after fractionation by gel filtration, the LBC shift is typically slightly stimulated by the inclusion of non-specific serum.

If the LBC binding to dHS1 is important for boundary function in the dHS1+HS3 combination, then it should be disrupted when the dHS1^mGAF^ fragment is used for the replacement instead of the wild type dHS1 fragment. Figure 5 shows that this is the case. Like the HS3 replacement alone, the dHS1^mGAF^+HS3 combination exhibited a strong GOF phenotype.

**Fig. 5.**
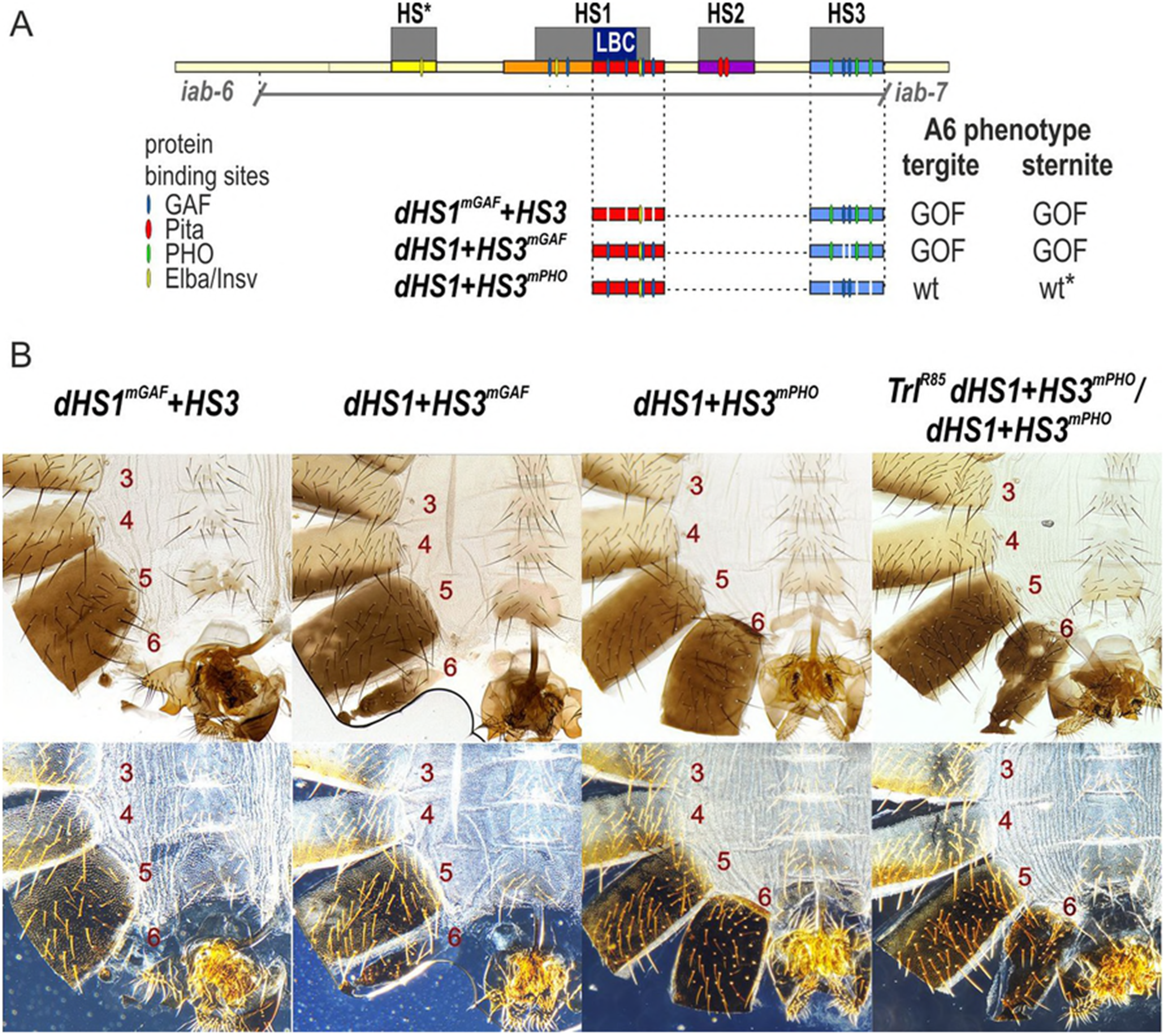
HS3 boundary activity requires the two GAGAG motifs but not the three Pho recognition sequences. (A) Map of *Fab-7* region showing the four hypersensitive sites, HS*, HS1, HS2, and HS3 and the locations of recognition motifs for proteins known to be associated with *Fab-7*. Replacement fragments are shown below the map with a summary of their cuticle (tergite and sternite) phenotypes. (B) Bright field (top) and dark field (bottom) images of cuticles prepared from *dHS1+HS3*^*mGAF*^, *dHS1+HS3*^*mPho*^, *Trl*^*R85*^ *dHS1+HS3*^*mPho*^, / *dHS1+HS3*^*mPho*^, and *dHS1*^*mGAF*^*+HS3* male flies. As described in text, most *dHS1+HS3*^*mPho*^ flies have a wild type phenotype, while the other mutant replacements typically exhibit strong GOF transformations. While nearly all *dHS1+HS3*^*mPho*^ males are wild type, reducing the dose of the *Trl* gene in half induces a GOF transformation. The sternite is usually absent while the tergite had an irregular shape and is reduced in size. wt* -- minor deviations in phenotype.

### The GAGA motifs in HS3 are required for boundary function while the Pho motifs are dispensable

A number of different mechanisms could potentially explain how HS3 contributes to the boundary functions of the *Fab-7* hypersensitive regions HS*, HS1, and HS2. One intriguing possibility is that the PcG dependent silencing activity of the HS3 *iab-7* PRE is needed for the boundary activity of replacements that contain only HS1/dHS1 (or HS1 plus either HS* or HS2). The HS3 *iab-7* PRE has two GAF recognition sequences (GAGAG) and three recognition sequences for the zinc finger protein Pleiohomeotic (Pho) [45]. Like many other *Drosophila* PREs [45,46], these DNA binding motifs are important for the PcG dependent silencing activity of the *iab-7* PRE. Mutations in either the GAGAG or Pho sequences compromise the silencing activity of the *iab-7* PRE in *mini-white* transgene assays [47]. Moreover, consistent with a role for the GAF (Trl) and Pho proteins in silencing, mutations in the *Trl* and *pho* genes suppress the silencing activity of the HS3 *iab-7* PRE [33,48].

If the PcG dependent silencing activity of the *iab-7* PRE is required to complement the boundary defects of HS1, then mutations in either the HS3 GAGA or Pho sequences should abrogate the boundary activity of the dHS1+HS3 combination. To test this prediction, we generated *dHS1+HS3*^*mGAF*^ and *dHS1+HS3*^*mPho*^ replacements. As expected, *dHS1+HS3*^*mGAF*^ lacks boundary activity, and flies carrying this replacement exhibit a strong GOF transformation of A6 (PS11) (Figure 5). However, contrary to our predictions, most *dHS1+HS3*^*mPho*^ flies are fully wild type, and exhibit no evidence of either GOF or LOF transformation. As was reported by Mihaly et al [29] for HS3 deletions, a small percentage (∼2-5%) of the male *dHS1+HS3*^*mPho*^ flies have a weak GOF transformation. In these flies the size of the tergite is reduced and/or the sternite is misshapen.

### The LBC binds to HS3 and binding depends upon two GAGA motifs

The fact that mutations in the GAGA motifs disrupt the boundary activity of HS3, while those in the Pho sites do not would argue that the PcG dependent silencing activity of HS3 is probably not responsible for its ability to complement HS1. Instead, it would appear that HS3 boundary function is separable from PRE activity. With aim of identifying factors contributing to HS3 boundary activity, we used three overlapping probes spanning HS3 (227 bp) for EMSA experiments with embryonic nuclear extracts. The EMSA experiment in Figure S5 shows that each probe generates several shifts. Though the identity of most of these shifts is unknown, the prominent slowly migrating shift observed with probe 2 resembles the shift generated by the LBC.

To determine if this slowly migrating shift corresponds to the LBC, we used a 200 bp fragment containing most of HS3, rather than the shorter probe 2. Like probe 2, the larger probe generates an LBC-like shift (Figure 4B). Two experiments indicate that this HS3 shift corresponds to the LBC. The first is a competition experiment with two different DNA fragments known to bind the LBC, *Fab-7* G3+4 and CES *roX2*. As could be predicted, the HS3 shift is competed by itself and also by excess unlabeled G3+4 and *roX2*. In the second experiment, we used peak LBC fractions from a gel filtration column, plus a control fraction that lacks LBC activity for antibody “supershift” experiments. Previous studies showed that the LBC shift is sensitive to antibodies direct against Clamp, GAF, and Mod(mdg4). Figure 4D shows that LBC binding to HS3 in the peak gel filtration fractions 45 and 47 is inhibited by Clamp antibody, while GAF and Mod(mdg4) antibodies generate a supershift.

These experiments indicate that like dHS1, the LBC binds to the full length HS3 sequence *in vitro*. This finding suggests a plausible mechanistic explanation for why HS3 contributes to *Fab-7* boundary and is able to reconstitute boundary activity when combined with dHS1. If this explanation were correct, we would expect that LBC binding to HS3 should require the GAGA motifs but not the Pho binding sequences. This is indeed the case. Figure 4C shows that mutations in the HS3 GAGA motifs (HS3^mGAF^) abrogate the LBC shift, while mutations in the Pho binding sequence (HS3^mPho^) have no effect. The requirement for the GAGA motifs, but not the Pho binding sequences is confirmed by competition experiments (Figure 4B) with mutant HS3 DNAs. HS3^mPho^ competes as well as the wild type HS3 for LBC binding, while HS3^mGAGA^ is a poor competitor.

### Protein occupancy is altered by mutations that impair function

To extend this analysis, we used chromatin immunoprecipitation (ChIP) experiments to compare GAF association with wild type and mutant versions of dHS1+HS3 in embryos and pupae. In the wild type *dHS1+HS3* replacement, GAF is found associated with both dHS1 and HS3 in embryos and pupae (Figure 6). As would be predicted from the loss of LBC binding to dHS1^mGAF^ DNA in nuclear extracts, GAF association with dHS1 in the *dHS1*^*mGAF*^ *+HS3* replacement is substantially reduced. Interestingly, GAF association with HS3 is also reduced in the *dHS1*^*mGAF*^*+HS3* compared to the wild type *dHS1+HS3* replacement. This secondary effect is seen not only in embryos, but also in pupae (Figure 6).

**Fig. 6.**
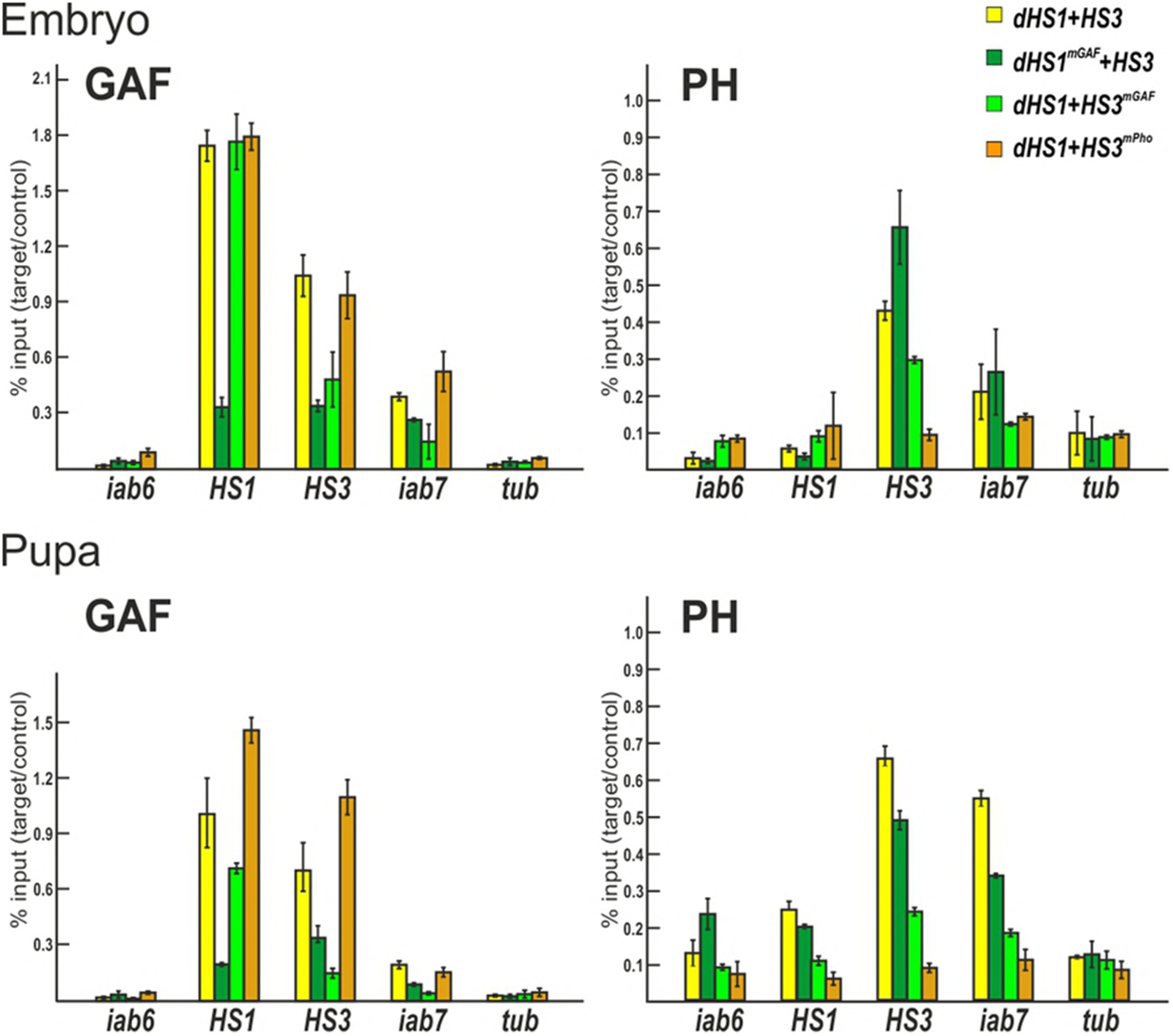
ChIPs with GAF and Ph antibodies at sites across the *Fab-7* region. Binding of GAF and Ph across the *Fab-7* region (*iab-6*, HS1, HS3, and *iab-7*) in different *Fab-7* replacements in embryos and pupae. The results of ChIPs are presented as a percentage of the input DNA normalized to a positive genomic site (*Hsp70* region – for GAF binding, PRE of *engrailed -* for Ph binding. Negative control is the *γTub37C* (*tub*) gene. Error bars indicate standard deviations of triplicate PCR measurements from three independent biological samples of chromatin.

Our EMSA experiments predict that GAF association with HS3 will be disrupted by mutations in the GAGA sequences, but not the Pho sequences. This expectation is correct. GAF association with HS3 is reduced in the *dHS1+HS3*^*mGAF*^ replacement while mutations in the Pho site (*dHS1+HS3*^*mPho*^) have no effect (Figures 4C and 6).

We also compared the dHS1/HS3 association of the Polycomb Repressive Complex 1 (PRC1) protein Polyhomeotic (Ph) in the different dHS1+HS3 replacements. As would be predicted from our previous studies on the silencing activity of HS3 in transgene assays, mutations in either the GAGA or Pho sequences reduce Ph-HS3 association (Figure 6). Ph also binds to a nearby sequence in the *iab-7 cis*-regulatory domain and, to a lesser extent, to HS1. In both cases, this association is reduced by mutations in the HS3 GAGA and Pho sequences. By contrast, mutations in the dHS1 GAGA sequences (dHS1^mGAF^) have limited effects on Ph association in HS3 or *iab-7*.

Our mutant replacement experiments showed that *dHS1+HS3*^*mPho*^ has nearly full boundary activity, while boundary activity is compromised in *dHS1+HS3*^*mGAF*^. These findings, together with our *in vitro* experiments on the LBC, would predict that the boundary activity of *the dHS1+HS3*^*mPho*^ replacement should depend on the GAF protein. If this is case, boundary activity might be sensitive to the dose of the gene encoding the GAF protein, *Trl*. To test this possibility we recombined the *dHS1+HS3*^*mPho*^ replacement with a *Trl* null mutation, *Trl*^*R85*^. The GOF transformations evident in *dHS1+HS3*^*mPho*^/*Trl*^*R85*^ *dHS1+HS3*^*mPho*^ flies show that the boundary activity of this replacement is compromised by a reduction in the dose of the *Trl* gene (Figure 5B).

## Discussion

Previous functional studies indicated that the *Fab-7* region of BX-C is subdivided into two seemingly distinct elements [29]. One of these elements spans the distal nuclease hypersensitive site HS3 and corresponds to a PRE for the *iab-7* regulatory domain. The other element spans the three proximal nuclease hypersensitive sites, HS*, HS1, and HS2. This element corresponds to the *Fab-7* boundary. In each case, the functional assignment was based on a combination of transgene assays and analysis of deletions in the *Fab-7* region generated by excision of a P-element insertion located between HS2 and HS3. This functional analysis has recently been extended with an *attP* replacement platform in which the entire region has been deleted [34]. This *attP* platform makes it possible to systematically test the functional properties of different *Fab-7* sequences in their native context. Using this platform, we recently found that hypersensitive site HS1 was both necessary and sufficient for wild type *Fab-7* boundary activity in a context in which the *iab-7* PRE, HS3, was present.

Since boundary activity in transgene assays required a sequence spanning HS*, HS1, and HS2, this finding suggested that the demands for full activity are considerably less stringent in the native context than in enhancer blocking assays. To confirm this conclusion, we tested the HS1 by itself. Unexpectedly, it is not sufficient for wild type boundary activity. Likewise, combinations of HS1 with either HS* or HS2 have only partial, tissue-specific boundary activity. In both combinations, the A6 sternite in males is absent, indicative of a PS11➔PS12 GOF transformation. For the HS1+HS2 combination, the A6 tergite is wild type, while, for the HS*+HS1 combination, the tergite displays weak to moderate GOF phenotypes. The only combination of these *Fab-7* sequences that has nearly complete boundary activity in the native context is HS*+HS1+HS2.

Like HS1 alone, the impaired boundary function of HS*+HS1 and HS1+HS2 can be rescued by the addition of HS3. These findings argue that HS3 must be able to contribute to *Fab-7* boundary function. Two different models could potentially explain the boundary activity of HS3. In the first, its boundary function would depend on the ability of HS3 to recruit PcG proteins and induce silencing. In the second, boundary function would reflect a PRE associated activity that is independent of PcG recruitment and silencing. For example, since PcG silencing is facilitated by pairing interactions between PRE containing transgene inserts on each homolog, the *iab-7* PRE, HS3, might have a chromosome architectural activity just like the classical boundaries [48-52]. A number of lines of evidence are consistent with this second model.

We compared the factors binding to HS1 and HS3 sequences using EMSA experiments with embryonic nuclear extracts. While probes spanning HS3 gave multiple shifts, the most prominent HS3 shift corresponds to a HS1 boundary factor, the LBC. This conclusion is supported by several observations. First, the HS3 LBC shift is competed by two DNA sequences, GAGA3+4 (*Fab-7*) and *roX2*, which are known to bind the LBC. Second, as observed for other LBC recognition sequences, antibodies against GAF, Clamp, and Mod(mdg4) either generate a supershift or interfere with binding to the HS3 sequence. Third, the peak LBC fractions after gel filtration of embryonic nuclear extracts have an apparent molecular weight of ∼1,000 kD. These peak LBC fractions generate a shift with the HS3 probe and were used for the antibody “supershift” experiments.

LBC binding to the *Fab-7* probes GAGA3, GAGA4, and dHS1 and to three X-linked CES depends upon GAGAG motifs (or GA rich sequences). This is also true for HS3. Mutations in the two HS3 GAGA motifs substantially reduce the yield of the HS3 LBC shift. Consistent with this finding, competition experiments indicate that the HS3^mGAF^ is a poor competitor for LBC binding to the wild type HS3 probe. While the HS3 GAGAG motifs are required for LBC binding, the two Pho sites are not.

Previous studies on the HS3 *iab-7* PRE indicate that like other fly PREs, it requires the GAF and Pho proteins for its silencing (and pairing-sensitive) activity [33,47,48]. Mutations in the two GAGAG motifs and in the two Pho recognition sequences disrupt silencing activity. Pho interacts with Sfmbt and is directly involved in the recruitment of PRC1 to PREs [53-56]. In vitro, GAF facilitates Pho binding to a chromatinized template [57].

If PcG silencing activity is critical for HS3 boundary activity, then boundary activity should be eliminated by mutations in either the GAGAG motifs or the Pho binding sites. In contrast, if LBC binding to HS3 is important, then mutations in the GAGAG motifs should disrupt the boundary function of the dHS1+HS3 replacement, while mutations in the HS3 Pho binding sequences should not. Consistent with the expectations of the second model, we found that the HS3 GAGAG motifs are important for boundary function, while the Pho binding sequences are not.

This distinction is also reflected in the protein occupancy of wild type and mutant HS1+HS3 replacements. In the wild type replacement, both GAF and Pho are associated with HS3. As would be predicted from the effects of GAGAG mutations on the PcG silencing activity of HS3 in transgene assays, the levels of both GAF and the Polycomb protein Ph are reduced in the HS1+HS3^mGAF^ replacement. In contrast, mutations in the HS3 Pho sites have no effect on GAF occupancy, while they reduce Ph occupancy. A prediction that follows from these findings is that the GAF protein is important for the boundary activity of dHS1+HS3^mPho^ replacement. Consistent with this prediction, dHS1+HS3^mPho^ boundary function is compromised when the flies are heterozygous for a mutation in *Trl*.

While the findings reported here support the idea that the boundary activity of both dHS1 and HS3 is mediated, at least in part, by the LBC, many questions remain. For example, why does HS3 *iab-7* PRE have PcG silencing activity while HS1 doesn’t? Likewise, why are the X-chromosome CES able to recruit the Msl dosage compensation complexes? One idea is that the silencing activities of the HS3 *iab-7* PRE and the dosage compensation functions of the CES depend upon the association of functionally specialized ancillary factors with a platform that is provided by LBC binding. This idea would be consistent with our findings. Both the silencing and boundary activities of the HS3 *iab-7* PRE depend upon GAF, while only the silencing activity depends upon Pho (which in other PREs is thought to function in the recruitment of PRC1). That functionally specialized factors might associate with different LBC recognition elements would also fit with our gel filtration experiments. We found that the LBC shifts in nuclear extracts (with *Fab-7* and several CES probes) migrate more slowly than the shifts observed after gel filtration. This difference in mobility suggests that there are factors that associate with the LBC:DNA complex in nuclear extracts that are not integral components of the LBC, and thus don’t co-fractionate with the LBC during gel filtration. These factors could contribute not only to the PcG silencing or MSL recruitment activities of individual LBC recognition elements, but also to the boundary activity of elements like *Fab-7* dHS1.

## Materials and Methods

### Generation of the *Fab-7*^*attP50*^ replacement lines

The strategy of the creation of the *Fab-7*^*attP50*^ landing platform and generation of the *Fab-7* replacement lines is described in detail in [34,36]. DNA fragments used for the replacement experiments were generated by PCR amplification and verified by sequencing. The sequences of the used fragments are shown in the Supporting Table S1.

### Cuticle preparations

Adult abdominal cuticles of homozygous eclosed 3-4 day old flies were prepared essentially as described in [36]and mounted in Hoyer’s solution. Photographs in the bright or dark field were taken on the Nikon SMZ18 stereomicroscope using Nikon DS-Ri2 digital camera, processed with ImageJ 1.50c4 and Fiji bundle 2.0.0-rc-46, and assembled using Impress of LibreOffice 5.3.7.2.

### Nuclear extracts

Nuclear extracts from 6- to 18-h embryos were prepared as described previously (Aoki et al., 2008) with small modifications. Embryos from Oregon R were collected from apple juice plates and aged 10 h at room temperature. The extraction was completed with the final concentration of KCl at 360 mM. Fractionation of the nuclear extracts derived from 6- to 22-h embryos was performed by size exclusion chromatography using Superose 6 10/330 GL column (GE Healthcare). Molecular mass markers ranging from 1,350 to 670,000 Da (Bio-Rad) were used as gel filtration standards.

### Electrophoretic mobility shift assay (EMSA)

Electrophoretic mobility shift assays were performed using γ-^32^P-labeled DNA probes under conditions described previously (Wolle et al., 2015). Probes for EMSA were obtained by PCR, purified on agarose– 1XTris-acetate-EDTA (TAE) gel followed by phenol/chloroform extraction. Probe sequences are listed in Supporting Table S1. Purified DNA probes (1 picomole) were 5’ end labeled with [γ-^32^P]ATP (MP Biomedicals/ Perkin Elmer) using T4 polynucleotide kinase (New England Biolabs) in a 50 μl total reaction volume at 37°C for 1 h. Samples were run through columns packed with Sephadex G-50 fine gel (Amersham Biosciences) to separate free ATP from the labeled probes. The volume of the sample eluted from the column was adjusted to 100 μl using deionized water so that the final concentration of the probe was 10 fmol/ μl. Binding reactions were performed in a 20 μl volume using the conditions described previously [34] except for the concentration of the non-specific competitor poly(dA-dT):poly(dA-dT) in the binding reaction. The final concentration of poly(dA-dT):poly(dA-dT) was varied between 0.1 and 0.25 mg/ml depending on the DNA probe used. 1 μl of nuclear extract (corresponding to about 20 ng of protein) or an equal volume of 360 mM nuclear extraction buffer (for negative control) was used. 2 or 3 μl of nuclear extract was used when indicated. In some reactions, unlabeled competitor DNA was included so that the final concentration of the competitor would be in 25- to 100-fold excess. The reaction mixtures containing the γ-^32^P-labeled DNA probes were incubated for 30 min at room temperature.

For supershift experiments, pre-immune rabbit serum or antibodies against different proteins were pre-incubated in the reaction mixtures described above with the nuclear extract or gel column fractions for 30 min at room temperature to allow the protein-antibody association, followed by an incubation with ^32^P-labeled DNA probes for 30 min at room temperature. Either 4 μl of rabbit polyclonal anti-CLAMP antibody [58], 1 μl of rabbit polyclonal antibodies against GAF and Mod(mdg4) was used.

Binding reactions were electrophoresed using the conditions described previously [34]. The gels were run at 180 V for 3 to 4 h at 4°C, dried, and imaged using a Typhoon 9410 scanner and Image Gauge software or X-ray film.

### Antibodies

ChIP antibodies against GAF (full length) were raised in rats and purified from the sera by ammonium sulfate fractionation followed by affinity purification on the CNBr-activated Sepharose (GE Healthcare, United States) according to standard protocols. Anti-Ph rabbit antibodies used in ChIP experiments were a gift from Maxim Erokhin. EMSA antibodies against GAF were obtained as gift from Carl Wu and David Gilmour, against Mod(mdg4) – from Anton Golovnin and Elissa Lei.

### Chromatin Immunoprecipitation

Chromatin for the subsequent immunoprecipitations was prepared from 12-24 h embryos and mid-late pupae as described in [39,59]. Aliquots of chromatin were incubated with antibodies against GAF (1:200), and Ph (1:500), or with nonspecific rat or rabbit IgG (control). At least three independent biological replicates were made for each chromatin sample. The results of the ChIP experiments are presented as a percentage of the input genomic DNA after triplicate PCR measurements and normalized to a positive genomic site for the appropriate protein, in order to correctly compare different transgenic lines with each other. The *γTub37C* coding region (devoid of binding sites for the tested proteins) was used as negative control; *Hsp70* region was used as positive control for GAF binding, PRE of *engrailed* was used as positive control for Ph binding. The sequences of used primers are presented in Supporting Table S1.

## Acknowledgments

We thank Farhod Hasanov and Aleksander Parshikov for fly injections. We would especially like to thank François Karch for the use of the *Fab-7 attP* replacement platform. This study was supported by the Russian Science Foundation, project no. 14-24-00166 (to P.G.) and by NIH to PS (R01GM043432 and R35GM13014). This study was performed using the equipment of the IGB RAS facilities supported by the Ministry of Science and Education of the Russian Federation.

## Supporting Figures

**Fig. S1 Variations in morphology of the abdominal segments of the *HS3* males.** Brightfield and darkfield images of male cuticles, as indicated. *Fab-7*^*attP*^: A6 is absent, indicating that PS11 is transformed into a duplicate copy of PS12. *HS3*: Two different classes of phenotypes are observed. The first class (I) resembles the GOF transformation of the starting *Fab-7*^*attP*^ replacement platform. The second class (II) has a small residual tergite that (based on trichome hairs) appears to have an appropriate A6 (PS11) identity. Sternites are not observed in either class. *HS1*: Three classes, I, II, and III, are observed. These classes differ in the size of the tergite. Class II is the most frequent.

**Fig. S2 Variations in morphology of the abdominal segments of *HS*+HS1* and *HS*+HS1+HS2* males.** Brightfield and darkfield images of male cuticles, as indicated. *HS*+HS1*: The cuticular phenotypes fall into three different classes depending on the size of the A6 tergite. In the most frequent class, class I, the tergite is significantly reduced in size and misshapen. There is a modest reduction in the size of the tergite in class II, while in class III, which is the least frequent, there is only a slight reduction in the size of the tergite compared to wild type. In these flies, trichome pattern in the A6 tergite resembles that in wild type, suggesting that surviving histoblasts that give rise to the dorsal cuticle are properly specified. In all *HS*+HS1* male flies the A6 sternite is missing. *HS*+HS1+HS2*: Three classes of cuticular phenotypes are observed. In the most frequent class, class III, the size of tergite is close to that in wild type, though sometimes the edges of the tergite are irregular. The sternite is present, but typically misshapen. Flies in the next most frequent class, class II, lack a sternite, while their tergite resembles that of class I. Finally, in class III, the tergite is noticeably reduced in size, while the sternite is misshapen.

**Fig. S3 Variations in morphology of the abdominal segments of the *pHS1+HS3* males.** Two roughly equal classes of phenotypes are observed for the *pHS1+HS3* replacement. In class I, A6 is transformed into a duplicate copy of A7, and is absent. In class II, the transformation is not complete, and a small residual A6 tergite is observed.

**Fig. S4 LBC binds preferentially to larger fragments.** Nuclear extracts prepared from 6-18 hr embryos were used for EMSA experiments: (-) no extract, (+) with extract. Comparison of LBC binding to probes spanning just GAGA3 (G3) or GAGA4 (G4) to probes spanning both GAGA3 and GAGA4 (G3+4). (A) EMSAs of G3, G4, and G3+4. (B) EMSAs of G3 and G3+4 with increasing amount of extract (1 μl, 2 μl, 3 μl). (C) Competition experiments with probe G3+G4 and excess cold G3+G4 or G3+LacZ (left to right: 100x, 75x, 50x, 25x, and10x).

**Fig. S5 LBC binding to HS3.** Nuclear extracts prepared from 6-18 hr embryos were used for EMSA experiments with three overlapping HS3 probes: Probe #1, 100 bp from proximal side of HS3. Probe #2, 100 bp probe from center of HS3. Probe #3, 88 bp probe from distal side of HS3. * – unique shifts; arrows – shifts observed with two or more probes.

**Supporting Table S1. The list of oligonucleotides and DNA fragments.**

